# Multimetric MRI Captures Early Response and Acquired Resistance of Pancreatic Cancer to KRAS Inhibitor Therapy

**DOI:** 10.1101/2024.11.22.624844

**Authors:** Mamta Gupta, Hoon Choi, Samantha B Kemp, Emma E Furth, Stephen Pickup, Cynthia Clendenin, Margo Orlen, Mark Rosen, Fang Liu, Quy Cao, Ben Z. Stanger, Rong Zhou

**Author notes:** Corresponding author: Rong Zhou, PhD, Department of Radiology, University of Pennsylvania, 198 John Morgan, Philadelphia, PA 19104.

## Abstract

**Purpose:** In pancreatic ductal adenocarcinoma (PDAC), KRAS mutations drive both cancer cell growth and formation of a dense stroma. Small molecule KRAS inhibitors (KRASi) represent a promising new treatment hence clinical tools that can assess early response, detect resistance and/or predict prolonged survival are desirable to understand clinical biology of KRASi. We hypothesized that diffusion-weighted MRI (DWI) can detect cell death while dynamic contrast enhanced MRI (DCE) and magnetization transfer ratio (MTR) imaging are sensitive to tumor microenvironment changes, and these metrics shed insights into tumor size change induced by KRASi treatment.

**Experimental Design:** Multiple preclinical PDAC models including a genetically engineered mouse model (KPC) received MRTX1133, a KRASi specific for KRAS^G12D^ mutation. Quantitative imaging markers were corroborated with immunohistochemistry (IHC) analyses.

**Results:** Significant increase of tumor apparent diffusion coefficient (a DWI metric) was detected as early as 48h and persisted to Day7 after initiation of KRASi treatment and was strongly correlated with cell death and reduced cellularity, resulting in greatly prolonged median survival in treated mice. Capillary perfusion/permeability (a DCE metric) exhibited an inverse relationship with microvascular density. Distinct responses of KRAS^G12C^ *versus* KRAS^G12D^ tumors to MRTX1133 were captured by the MRI metrics corroborated with IHC. When tumors developed resistance to MRTX1133, the imaging marker values exhibited a reversal from those of responding tumors.

**Conclusions:** Multiparametric MRI provides **early** biological insights of cancer and stromal response to KRASi treatment and sets the stage for testing the utility of these clinically ready MRI methods in patients receiving KRASi therapy.

**Translational relevance:** Emerging small molecule KRAS inhibitors (KRASi) represent a new class of therapy for PDAC. Clinical tools that can provide early biological insights of KRASi therapy are desirable. In PDAC models, we examined a clinically ready imaging protocol that combines MRI-based tumor size, diffusion-weighted MRI (DWI), dynamic contrast enhanced MRI (DCE), and magnetization transfer ratio (MTR) for detection of early response as well as acquired resistance to MRTX1133, a KRASi being evaluated in clinical trials. Our data show that DWI and DCE metrics provided key insights of significant cell death and tumor microenvironment changes underlying tumor size regression as early as 48 hours after KRASi treatment initiation. These MRI metrics also captured resistance to KRASi developed over prolonged treatment. This study has high translational relevance by employing clinically applied MRI methods, an investigational new drug and a genetically engineered mouse model that recapitulates salient features of human PDAC.

## Introduction

The oncogene RAS is involved in 1/3 of human cancers, and among the 3 isoforms of RAS (KRAS, HRAS and NRAS), KRAS mutations account for a majority (85%) of RAS mutations and occur almost exclusively in pancreatic ductal adenocarcinoma (PDAC), where 98% patients carrying KRAS mutations at glycine 12 position (KRAS^G12X^) (1). The mutant RAS protein exists predominantly in an active, GTP-bound state, leading to hyperactive signaling of cell proliferation and survival (2). KRAS mutation also drives the formation of a desmoplastic stroma, which restricts drug delivery and an immune suppressive tumor microenvironment (3–5). Recent breakthroughs in the development of small molecule RAS inhibitors (RASi) have led to FDA approval of two KRAS^G12C^ inhibitors, Adagrasib and Sotorasib, for lung cancer treatment. Furthermore, RASi targeting G12D, G12V, G13X and Q61X mutations and wild type RAS are being evaluated in various phases of clinical trials (6–10). Together with other RAS inhibiting modalities (11,12), this new class of RAS inhibitors hold the potential to transform the treatment of RAS-driven cancers especially PDAC, which remains the third-leading cause of cancer-related deaths.

To assess responses of PDAC tumors to KRAS inhibitor (KRASi) therapy in the clinic, we expect RECIST-based tumor size (13) to provide an objective metric of tumor regression observed in earlier studies (9,14). However, insights into tumor size reduction are desirable. Specifically, cancer cell death can result from the blockade of ERK signaling upon engagement of KRASi to its target (15), meanwhile, a mechanism that KRAS inhibition induces T cells mediated cancer cell killing is plausible (16). Indeed, significant apoptosis of PDAC cells was detected in tumor specimens at early stage of KRASi treatment (14) hence clinical tools that can detect early cell death would be useful to confirm the drug-target engagement. Besides cell death, stromal changes induced by KRASi also contribute to tumor size regression and have a great impact on tumor microenvironment. Knowledge of microvascular perfusion and permeability is important since these properties influence lymphocytes attachment to capillary lumen and extravasation into tumor parenchyma (17–19).

Based on these considerations, we evaluated clinically ready MRI modalities including diffusion weighted MRI (DWI), dynamic contrast enhanced MRI (DCE) and magnetization transfer ratio (MTR) in addition to standard of care tumor size measurement in the setting of KRASi therapy. While DWI markers associated with other diffusion regimens (20,21) have been applied in the clinic studies, apparent diffusion coefficient (ADC) derived from Gaussian diffusion model appears to be most robust, capable of early detection of tumor response and prediction of treatment outcomes in many clinical studies - reviews/ editorial (22–26). ADC quantifies the *hindered* diffusion of water molecules in *extracellular space* of tumor tissues due to the presence of cell membrane and extracellular matrix components, making ADC most sensitive to *extracellular space volume fraction and geometry* (27). For example, significant cancer cell death induced by chemotherapy and radiotherapy respectively increased the tumor ADC compared to pretreatment values, while tumor ADC was shown to positively correlate with apoptosis and inversely correlate with tumor cellularity (28,29). We <u>hypothesized</u> that in tumors responding to KRASi, an increase of ADC early after KRASi treatment is indicative of cell death hence confirming drug-target engagement.

DCE offers a tool for evaluating microvascular function (perfusion and permeability) *in vivo*, and like ADC, the utility of DCE has been established in clinical studies of cancer therapies specifically for assessing their anti-angiogenic and /or anti-vascular effects (30). K^trans^, a DCE metric related to capillary perfusion and permeability, can detect early responses and predict complete pathological response to chemotherapy (31) and reveal vascular effect mediated by stroma directed drug against PDAC (32). We <u>hypothesized</u> that the DCE metrics *K*^*trans*^ and extracellular & extravascular volume fraction (V_e_) can detect KRASi-induced changes in microvascular functional features.

MTR imaging is sensitive to the relative volume fractions of bound versus free water and the exchange rate between the two. Increased collagen deposition associated with pathological conditions such as fibrosis provides increased surface area for water molecules to bind, hence is associated with increased MTR (33,34).

We therefore <u>hypothesized</u> that MTR is sensitive to changes in matrix collagen induced by KRASi treatment. Recently, however, changes in MTR were related to alterations in cellularity in a study of bone marrow malignancy (35).

Taken together, tumor size, DWI, DCE and MTR constitute a *multiparametric MRI* (mpMRI) tool (described in **Supplementary Figure S1**) that can potentially provide insights into clinical biology of KRASi therapy. To assess this approach, we conducted a preclinical trial of an investigational new drug, MRTX1133, a KRASi specific for KRAS^G12D^ mutation in several PDAC models including a genetically engineered mouse model of PDAC that features activating KRAS^G12D^ mutation and recapitulates key features of the human disease (36). mpMRI was applied on Day2 (48 h) and Day7 after initiation of KRASi therapy. Treatment was extended beyond Day7 in a survival study setting. We further compared mpMRI and IHC signatures of tumors at resistant *versus* responding stages which may be used for testing the combinational therapy aimed to overcome the resistance.

## Materials and Methods

MRTX1133 was provided by Mirati Therapeutics /Bristol Myers Squibb (Cambridge, MA) via an institutional material transfer agreement. All animal procedures were approved by the institutional animal care and use committee (IACUC) of the University of Pennsylvania. A list of abbreviations was provided in the Supplementary information.

### Mouse models and Study design

A genetically engineered KrasLSL^G12D^-Trp53LSL^R172H^-Pdx1-Cre (KPC) mouse model of PDAC (36) was bred in the Mouse Hospital of Pancreatic Cancer Research Center, University of Pennsylvania. Screening for tumors was done via weekly abdominal palpation starting at 11 weeks of age followed by ultrasound examination (Vevo 2100, VisualSonics, Toronto, ON, Canada) to estimate tumor size. Mice with a confirmed tumor mass (both sexes, 18–25 weeks old) were enrolled in the MRI study.

Two PDAC cell lines (4662-G12D and 4662-G12C) were obtained from the Stanger lab and used within 3 passages for this study. The 4662-G12D cells were derived from tumors from KPC tumors as we previously described (37). The 4662-G12C cell line was generated from 4662-G12D cells using CRISPR/Cas9 and retroviral transduction as we described (14). Cell lines were validated by RT-PCR to assess for Cre-mediated activation of the mutant *Kras* and *Trp53* alleles (37) and periodically tested for Mycoplasma using the MycoAlert Mycoplasma Detection Kit (Lonza, LT07-318). The 4662-12D and 4662-G12C cells were respectively inoculated in the flank of 7-week old C57BL/6 mice (Charles River) (RRID:SCR_003792) to grow subcutaneous (subQ) tumors. In separate mice, 4662-G12D cells were inoculated into the pancreas to grow orthotopic tumors as we described previously (32). The mice with tumor size below 50 mm^3^ at the baseline were excluded in the study.

Both male and female mice were used and were randomized to MRTX1133 or control (CNTRL) group. MRTX1133 (Mirati Therapeutics) at 30 mg/kg twice daily was formulated in vehicle (14) and injected intraperitoneally (ip), while CNTRL mice received 0.2 mL PBS ip twice daily.

As outlined in **Figure 1**, we first tested the utility of mpMRI to detect response on Day2 and Day7 of MRTX1133 treatment in KPC mice (Figure 1**A**). We then assessed the survival benefit of KRASi therapy and whether baseline mpMRI can predict the survival of treated mice (Figure 1**B**). Using isogenic model bearing KRAS^G12D^ and KRAS^G12C^ mutation, respectively, we then examined whether mpMRI can detect the specificity of MRTX1133 to KRAS^G12D^ mutation on Day2 (Figure 1**C**). We further monitored the KPC tumor through the time course over which the tumor developed resistance to MRTX1133 and we compared mpMRI metrics in resistant versus responding stages (Figure 1**D**).

**Figure 1:**
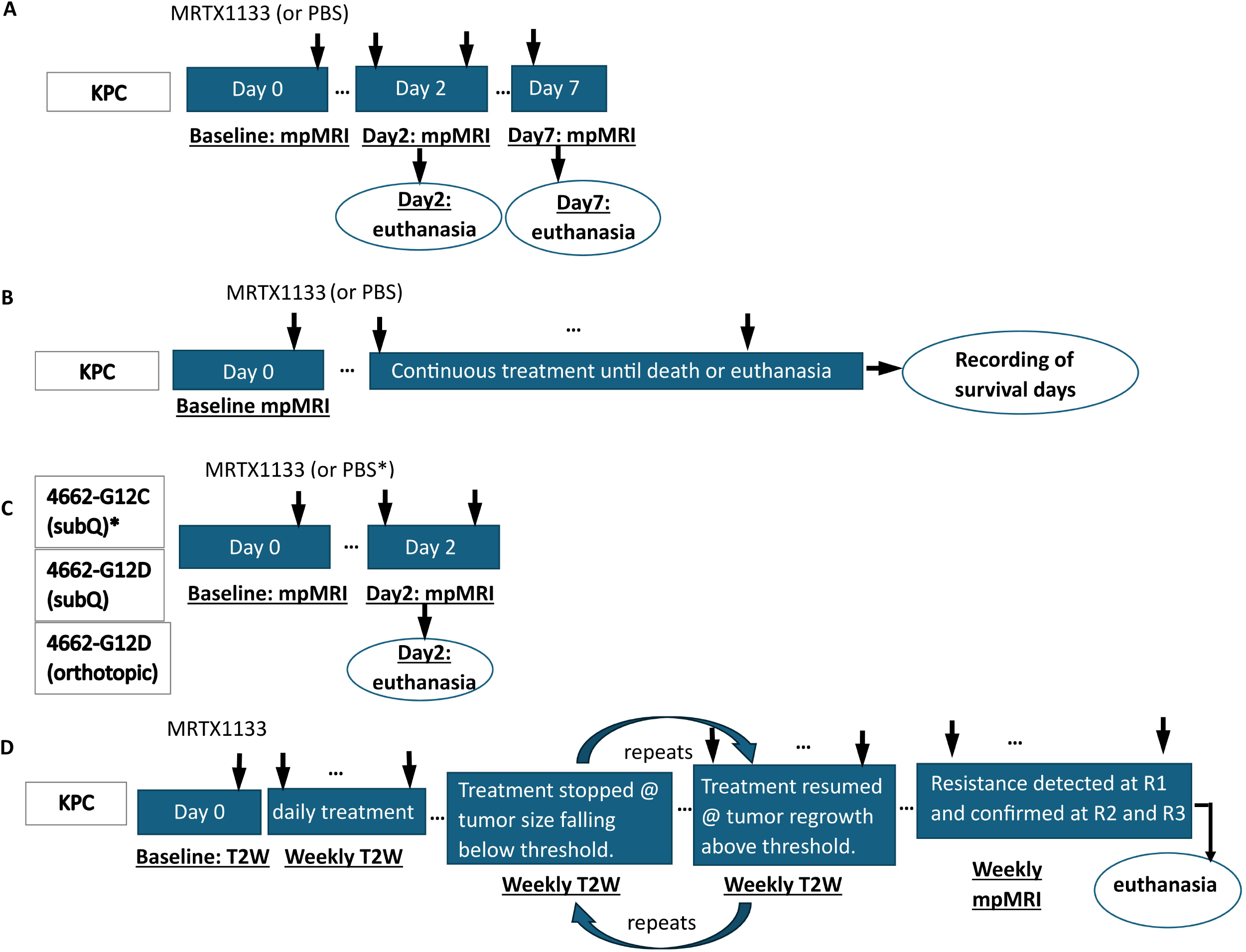
Experimental designs. **(A)** For detection of early and stable treatment effects of KRASi (MRTX1133), respectively, a genetically engineered mouse model (KPC) was studied at baseline, on Day2 and Day7 after treatment initiation by mpMRI followed by euthanasia for IHC analyses. **(B)** Survival study in KPC model with KRASi monotherapy or PBS (CNTRL). **(C)** Isogenic 4662-G12C and 4662-G12D PDAC models carrying KRAS^G12D^ or KRAS^G12C^ mutation, respectively were studied at baseline and Day2 after treatment initiation by mpMRI followed by euthanasia for IHC analyses. **(D)** Monitoring the development of resistance to KRASi monotherapy by weekly T2W MRI. Treatment was paused when the tumor was not measurable on MRI while weekly T2W MRI continued to capture tumor regrowth. Once the tumor size rose above radiographic threshold, treatment was resumed. This cycle was repeated until resistance was developed and captured by T2W MRI. Weekly mpMRI was performed on R1/2/3 stage. For all studies, KRASi or CNTRL treatment was initiated on Day0 after baseline MRI.

In survival study (Figure 1**B**), KPC mice were enrolled in MRTX1133 (n=20, tumor volume = 227±74 mm^3^) or CNTRL (n=20, tumor volume = 181±94 mm^3^) after baseline mpMRI and their survival durations were recorded. In case a mouse must be euthanized, the death was recorded for the next day. To monitor the development of resistance (Figure 1**D**), tumor size was measured weekly by MRI. If tumor size became radiographically undefined, the treatment was paused until it was measurable again. Resistance is defined as tumor size increases from prior week and resistant stage-1 (R1) was the first time such increase was detected; the mouse was monitored for 2 additional weeks (R2 and R3) to confirm no regression occurred. DCE data was collected only at R1 while other mpMRI metrics at all resistant stages.

### In vivo MRI studies

#### Animal preparation

General anesthesia was induced during preparation and maintained during MRI studies via breathing of 1-3% isoflurane in air through a nose cone. To facilitate contrast agent injection during DCE acquisition, a tail vein catheter was placed with an extension tubing preloaded with ProHance (Bracco Diagnostic, NJ, USA) diluted (50×) in PBS to 10 mM Gd. The animal was placed on a 3D printed bed, which was positioned in a 35 × 40 (ID x length) mm quadrature birdcage RF coil (M2M, Cleveland, OH, USA). During MRI, vital signs were monitored via a rectal temperature probe and pneumatic respiration pillow (SAII, Stonybrook, NY, USA), and core body temperature was maintained at 37 ± 0.2°C by warm air (32,38–40).

#### T2W, DWI, DCE and MTR acquisitions

MRI studies were performed on a 9.4 T Avance III console (Bruker, Berillica, MA, USA), equipped with 12 cm ID, 40 G/cm gradients. Following calibrations of the scanner and generation of scout images, an axially oriented contiguous series of T2W images spanning the entire tumor was generated using a TurboRare protocol (effective Echo time = 30ms, Repetition time = 1.8 s, echo train length = 8, matrix size = 64 × 64, slices = 16, Field Of View = 32 × 32mm, thickness = 0.5 mm, averages = 4, total acquisition time ~2 min). The image orientation and abdominal region coverage of the T2W scan were passed to DWI, DCE and MTR scans.

To mitigate respiratory motion-induced artifacts in DWI, a radial k-space sampling scheme (38,39) was employed with acquisition parameters of diffusion time (Δ) = 14 ms, diffusion gradient duration (δ) = 9 ms, b-value = 10, 535, 1070, 1479, and 2141 s/mm^2^, Echo time/Repetition time = 28.6/750 ms, matrix size = 64 × 403, averages = 1, bandwidth = 50 kHz with fat saturation and total acquisition time approx. 25 min. No respiration gating was employed while the mouse was breathing freely during acquisition. The detailed protocol can be found at protocols.io (dx.doi.org/10.17504/protocols.io.kxygxp6jol8j/v1).

To enable optimal spatial and temporal resolution of DCE, 3D stack-of-stars (SoS) sequence was applied to all acquisitions in the DCE protocol including B1 mapping by actual flip angle imaging (AFI), T1 mapping by variable flip angle (VFA) imaging and the DCE series by a golden-angle (111.25°) ordering scheme to obtain a uniform coverage of k-space as we described previously (40). Two minutes after starting the DCE series, 0.2 mL of diluted Prohance was injected over 10 s manually via the catheter. All components of the DCE protocol had matrix size = 64×201×16 and Field Of View = 32×32×8 mm^3^. The detailed protocol can be found at protocols.io (dx.doi.org/10.17504/protocols.io.81wgb6pnolpk/v1).

For MTR, a magnetization prepared 3D gradient echo was acquired with Field Of View= 32×32×8 mm, matrix size = 64×64×16, Echo time/Repetition time = 2.8 /5.7 ms, flip angle= 5°, averages = 4 with an 18 sec 2.5 µT saturation pulse applied at 4 kHz (*MT*_*on*_ image) and 250 kHz off-resonance (*Reference* image), respectively. Each saturation pulse was followed by acquisition of a full plane k-space data using centric phase encoding.

### MRI Data Analytics

**The tumor volume** was determined by manually tracing the tumor boundary on T2W images to generate tumor Region Of Interest, summing the areas from all Region Of Interest and multiplying the result by the slice thickness. DWI, DCE and MTR metrics were estimated over the same Region Of Interest.

#### DW-MRI metrics

Radial k-space data were reconstructed to images using custom python codes (*https://github.com/PennPancreaticCancerImagingResource/DWIProcessing*). The reconstructed images were subject to pixel-wise least squares fit to Equation [1] using all five b-values as described previously (39), where *S*_*0*_, *ADC*, and *KI* (kurtosis index) are the fitting parameters: *S*_*0*_ is the signal intensity in the absence of diffusion weighting and *KI* is a constant that accounts for contributions from water molecules undergoing slow hence non-gaussian diffusion to the observed signal *S(b)*.

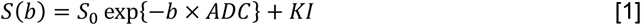

#### DCE-MRI metrics

Radial k-space data were reconstructed using k-space weighted image contrast (KWIC) method to obtain optimal spatial and temporal resolution for DCE images as we described previously (40). The shutter-speed kinetic model programmed in custom IDL codes (Harris Geospatial Solutions, Broomfield, CO, USA) was used for least-square fitting of the dynamic data to derive the DCE metrics: K^trans^ (rate constant of transferring unit volume of contrast agent from capillaries to interstitial space per minute), V_e_ (extracellular and extravascular volume fraction, %,) and τ_i_ (intracellular water life time in second) (41). Additional inputs to the fitting include T1 maps and a group arterial input function (AIF) derived from KPC mice (42).

#### MTR metrics

MTR value was calculated pixel-wise by equation [2] to generate MTR maps from tumor Region Of Interest:

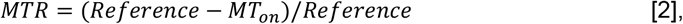

Where the *MT*_*on*_ and *Reference* image were acquired with and without saturation of motion-restricted water, respectively.

The range of MTR value (0-1) was divided into 25 equal width bins and the number of pixels in each bin was normalized to total number of pixels in the tumor. Since tumor MTR values were primarily distributed between 0-0.56 (1^st^ – 15^th^ bins) with less than 10 pixels per tumor whose MTR were greater than 0.56 hence 16^th^ - 20^th^ bins were combined into the 15^th^ bin. MTR histogram was generated from *MTR*_*Bin[x]*_, where x is the bin number (1, 2, …, 15).

For every MRI metric (ADC, KI, K^trans^, V_e_, MTR, MTR_Bin_), the median value is estimated from all pixels in the Region Of Interest of each tumor.

### Immunohistochemistry, Immunofluorescence and Collagen Assay

Immunohistochemistry (IHC) for Sirius Red, H&E and CD31 of formalin fixed paraffin embedded tumor sections were conducted by the Pathology Core of Children’s Hospital of Philadelphia (www.research.chop.edu/pathology/tools). Stained sections were scanned at ×40 magnification using Aperio ScanScope CS2 (Leica Biosystems Imaging, CA, USA) and uploaded to QuPath for analyses (43). For each section, tumor regions were defined, and hand drawn by a pathologist (EEF) specializing in GI cancer, who was blinded to the treatments. Sections stained with H&E, Sirius Red, and CD31 were used to determine viable cell density, % fibrosis, and microvascular density, respectively.

For immunofluorescence (IF) staining of pERK1/2, CK19, Ki-67, cleaved Caspase-3 (CC3), α-SMA, formalin fixed paraffin embedded slides were deparaffinized and rehydrated through a series of xylene and ethanol washes. Antigen retrieval was followed by blocking with 5% donkey serum for 1 hour at room temperature. Based on dilution schemes in **Supplementary Table S1**, primary antibodies were incubated with the slide for 1 hour at 4°C followed by 2× wash with PBS and incubation with secondary antibodies for 1 hour at room temperature. Nuclei were counterstained with DAPI (1:1000). Slides were mounted with aqua-poly mount and cover slipped. Images were captured using an Olympus IX71 inverted microscope with 4× and 20× objectives and a DP71 camera, and 3-5 magnified views per sample were analyzed using FIJI software.

To estimate collagen content, tumor tissue was homogenized in DI water (500 μL per 50 mg of tissue) in a homogenizer (Bertin Technology, Rockville, MD). The homogenate was transferred to a screw-cap tube to which 100 μL of 10 N NaOH was added, and the mixture was heated at 120°C for 1 hour. Afterwards, the tube was cooled followed by addition of 100 μL of 10 N HCl and centrifugation at 10,000 x g for 5 minutes. The supernatant was collected for estimation of collagen concentration according to the manufacturer’s instructions (ab222942, Abcam, Newark, DE).

### Statistical Analysis

For MRI and IHC metrics, the mean and standard deviation for each metric were calculated separately from the CNTRL and MRTX1133 treated group. Shapiro-Wilk tests were performed to assess whether the data followed a normal distribution. For datasets that passed the normality test, two-tailed t-tests were used to compare the metrics between CNTRL and MRTX1133. Nonparametric tests were used when normality assumptions are not met and specified in figure captions. Bootstrapping analysis was conducted to confirm the statistical significance of *MTR*_*Bin*_ values. MTR bins were categorized into 4 groups (i.e., 0-0.2, 0.2-0.4, 0.4-0.6, and 0.6-1), and the proportion of tumor pixels within each MTR range was calculated for baseline and Day2. Given the relatively small sample size (n=10), the upstrap method (44) was used to simulate a larger sample size and build the 95% confidence interval (CI) for the mean difference in proportions between baseline and post-treatment.

To assess the association between individual MRI metrics (tumor size, ADC, KI, MTR, K^trans^ and V_e_) and IHC markers of cancer, Pearson correlation analyses were performed. In addition, to assess whether the MRI metrics could be used to predict survival, regression analyses were conducted. Statistical analyses were conducted in GraphPad (RRID:SCR_002798) or R software, with α value set at 0.05. Due to the exploratory nature of the study, a formal power calculation was not conducted.

### Data availability

The raw data generated in this study will be available to share upon a request to the corresponding author.

## Results

### mpMRI captured the early (Day2) and stable (Day7) response to KRASi therapy

mpMRI study of therapeutic response on Day2 and Day7 enrolled separate KPC cohorts and tumor tissues were collected after imaging (**Figure 1A**). mpMRI and IHC results on Day2 are summarized in **Figure 2–3** and **Supplementary Figure S2 and S3** while those on Dday7 in **Figure 4** and **Supplementary Figure S4**.

**Figure 2.**
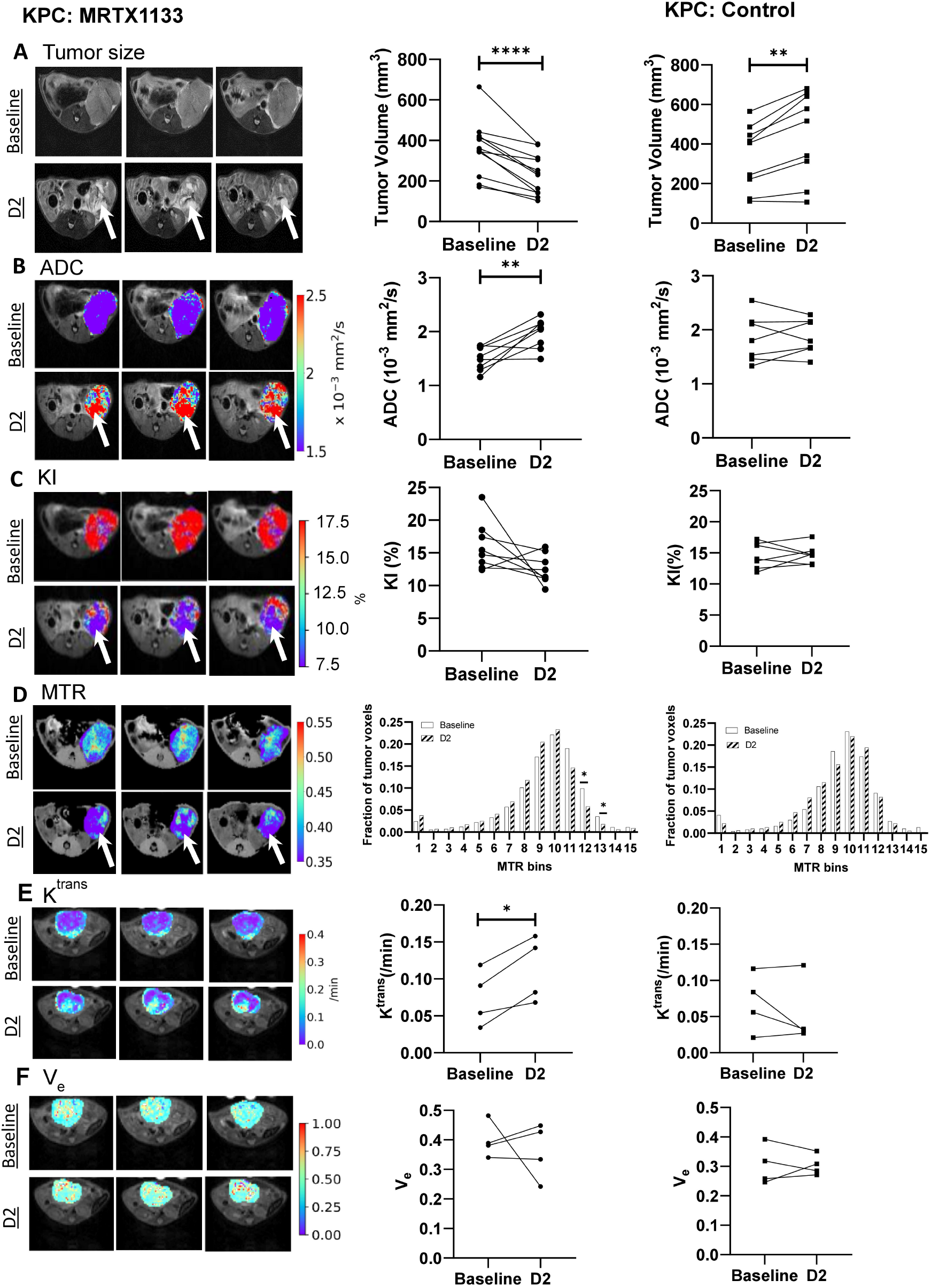
Day2 MRI metrics of KRASi-treated and CNTRL mice. (**A**)Tumor volume, (**B**) ADC, (**C**) KI, (**D**) MTR bin histogram, (**E**) K^trans^, and (**F**) V_e_ of the tumor from individual mice at baseline and Day2 (D2) are shown for treated and CNTRL mice. The tumor was defined on T2W MRI (yellow dotted line in **A**) while parametric maps of the tumor are overlaid on T2W images (**B-F**). Two-tail, paired t-test was conducted to compare Day2 vs. baseline. \**P* <0.05, ** *P* <0.01, *** *P* <0.001, **** *P* <0.0001.

**Figure 3:**
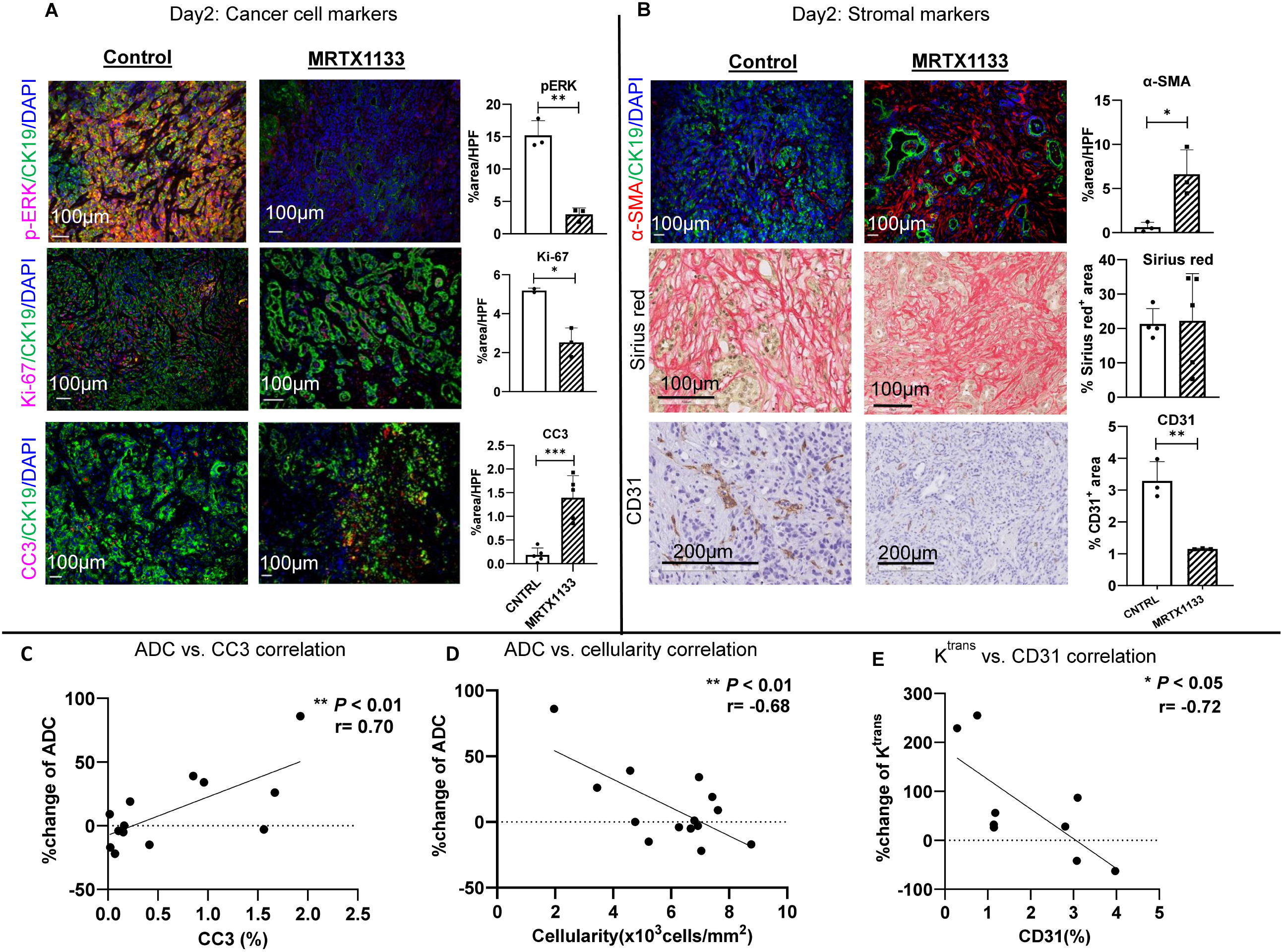
Day2 IHC markers of cancer cells and stroma and correlations with mpMRI markers. (**A**) p-ERK1/2 & CK19 double staining marked KRAS inhibition in cancer cells while proliferation and death of cancer cells are assessed by double staining of Ki67 & CK19 and CC3 & CK19, respectively. (**B**) Stromal markers include αSMA, Sirius red (for collagen) and CD31 for tumor microvascular density. αSMA-positive but CK19 negative cells were estimated in the plot. CD31-positive cells stained in brown color while Sirius red in red. (**C**) Correlation between ADC change vs. cancer cell death. (**D**) Inverse correlation between ADC change vs. cellularity estimated on the H&E-stained tumor sections. (**E**) Inverse correlation between K^trans^ vs. microvascular density. For C,D,E, mpMRI and IHC data from Day2 and Day7 are pooled. Two-tail, unpaired t-test was conducted to compare treated vs. CNTRL while Pearson correlation analysis was applied to C, D, E. \**P* <0.05, ** *P* <0.01, *** *P* <0.001.

**Figure 4:**
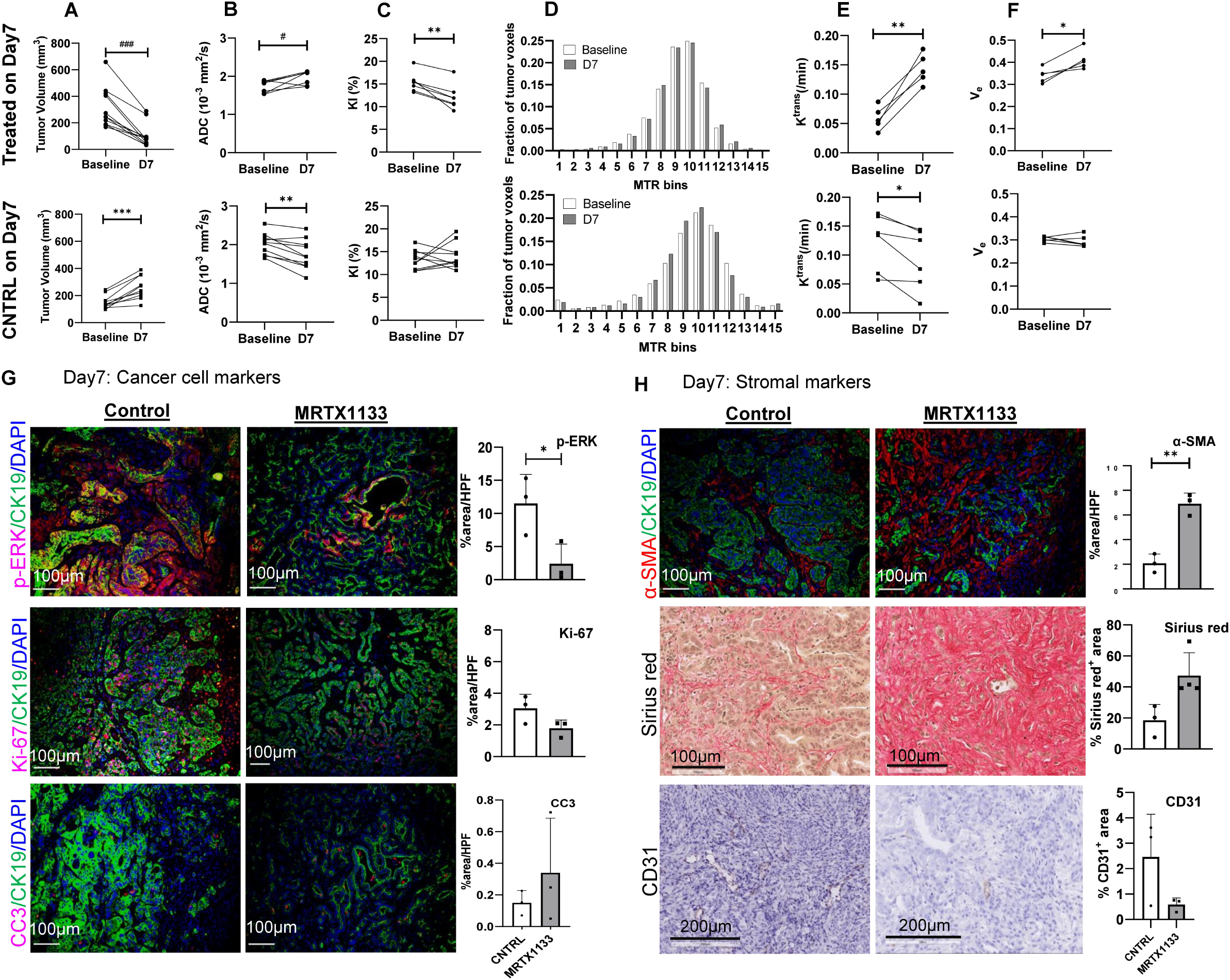
Day7 mpMRI and IHC of KRASi and CNTRL group (KPC mice). Tumor volume (**A**), ADC (**B**), KI (**C**), MTR (**D**), K^trans^ (**E**), and V_e_ (**F**) of the tumor from individual mice at baseline and Day7 (D7) are shown for treated and CNTRL mice. (**G**) KRAS inhibition in cancer cells is marked by p-ERK1/2 & CK19 double staining, proliferation and death of cancer cells are assessed by double staining of Ki67 & CK19 and CC3 & CK19, respectively. (**H**) Stromal markers of αSMA, Sirius red and CD31. αSMA-positive and CK19-negative cells were estimated in the plot. CD31-positive cells stained in brown color while Sirius Red in red. In (**A, B**), based on normality test outcome, the Mann-Whitney test was applied to tumor volume and ADC data with ^#^ *P* < 0.05, ^###^ *P* < 0.001. For remaining data, two-tail, paired or unpaired t-test was applied when appropriate: \**P* <0.05, ** *P* <0.01, *** *P* <0.001.

At baseline, tumors were readily delineated on T2W images (yellow dotted line, **Figure 2A**) and appeared homogenous. Parametric maps of tumor ADC, KI, MTR, K^trans^ and V_e_ in pseudo color overlaid on the T2W images were more homogeneous at baseline vs. Day2 (**Figure 2A-F**). MRTX1133 group had a 36% reduction of tumor size (from 359±133 mm^3^ to 231±99 mm^3^, n=12, *P* < 0.0001) in contrast to a 32% increase in CNTRL (from 335±164 to 443±220, n=9, *P* < 0.01). Meanwhile, a 31% increase in tumor ADC (from 1.50±0.22 x10^−3^ to 1.96±0.28 x10^−3^ mm^2^/s, n = 8, *P* < 0.01) in treated animals was highly significant as no change was observed in CNTRL (**Figure 2B**). To corroborate the mpMRI results, we evaluated cancer cell and stromal markers in tumor specimens collected after mpMRI (**Figure 3**): As a marker of KRAS/RAF/MEK pathway activity, a remarkable reduction in phospho-ERK (pErk1/2) level (15.2±2.3% vs. 3.0±1.0%/CNTRL, n=3 for both groups, *P* < 0.01, **Figure 3A**) confirmed the drug-target engagement. Consequently, significantly increased cancer cell death by CC3 staining (1.5±0.5% /Treated vs. 0.4±0.2% /CNTRL, *P* < 0.001, n = 5) and reduced cancer cell proliferation by Ki67 (2.5±0.7% /Treated vs. 5.2±0.1% /CNTRL, *P* < 0.05, n = 2-3) were observed and are consistent with an increased ADC since loss of membrane integrity and/or clearance of dead cells would increase diffusion path length of free water.

On Day7, pErk1/2 remained significantly suppressed, indicating persistent KRAS^G12D^ inhibition (**Figure 4G**), whereas increase in apoptosis (CC3) and reduction in proliferation (Ki-67) diminished compared to Day2, suggesting that extensive cell death occurred most intensively at early stage of KRASi therapy and waned overtime. However, the increase in ADC persisted to Day7 (*P* < 0.05, n = 7, **Figure 4B**), likely because clearance of dead cells sustained a low tumor cellularity. Combining Day2 and Day7 mpMRI and IHC data, a highly significant correlation between ADC and CC3 (*r* = 0.70, *P* < 0.01, **Figure 3C**) as well as an inverse correlation between ADC and cellularity (*P* < 0.01, *r* = −0.68, **Figure 3D**) were identified. These data provide a strong support that cell death and reduced cellularity underlie the increased ADC observed after KRASi therapy.

mpMRI also detected KRASi mediated changes in stroma and tumor microenvironment. First, a highly significant increase in DCE metric K^trans^ was observed on Day2 in treated animals (0.11±0.04 min^−1^ vs. baseline /0.07±0.04 min^−1^, n = 4, *P* < 0.05, **Figure 2E**), suggesting increased microvascular perfusion /permeability in response to KRASi therapy, whereas no changes were detected in CNTRL (**Figure 2E**). Increased K^trans^ persisted to Day7 (*P* < 0.01, n=5, **Figure 4E** upper). Paradoxically, microvascular density was significantly reduced on Day2 (1.2±0.0% /Treated vs. 3.3±1.2% /CNTRL, n = 3, *P* < 0.01, **Figure 3B**) with the same trend clearly shown on Day7 despite not reaching statistical significance (**Figure 4H**). Increased K^trans^ accompanied by reduced MVD on Day2 and Day7 led to a strong *inverse* correlation between *K*^*trans*^ *versus* microvascular density (*r* = −0.72, *P* < 0.05, **Figure 3E**).

Second, tumor collagen content was significantly increased on both Day2 and Day7 (**Supplementary Figure S2**), consistent with significantly increased αSMA staining (**Figure 3B** and **Figure 4H**) and the trend of increased Sirius Red staining on Day7 (**Figure 4H**). However, *no increase* in the mean MTR or MTR_bin_ values were detected on either timepoint (**Figure 2D** and **Figure 4D**), suggesting MTR parameters cannot mark the increased collagen content in KRASi therapy setting. Instead, *reductions* of MTR bin-12 and bin-13 (correspond to MTR ranging 0.4-0.56) were detected on Day2 compared to baseline in treated tumors (*P* < 0.05, n = 10, **Figure 2D**). This result was confirmed by the bootstrap analysis of MTR_bin_ (*P* < 0.05, n=10 for MTRX1133 and CNTRL group, respectively). Such reduction is consistent with cell death leading to reduced intracellular bound water pool.

The longer treatment period of 7 days led to more uniform treatment effects: the trend of reduction in KI detected on Day2 reached statistical significance on Day7 (*P* < 0.01, n = 7, **Figure 4C** upper panel) so did DCE metric, V_e_ (*P* < 0.05, n = 5, **Figure 4F** upper) consistent with deeper tumor regression (Day7/ 97±87 mm^3^ vs. Day2/ 231±99 mm^3^, *P* < 0.001, n = 12, **Figure 4A**). On Day7, CNTRL group exhibited 14% reduction in ADC (n= 10, *P* < 0.01, **Figure 4B** bottom) and 28% reduction in K^trans^ (n= 6, *P* < 0.05, **Figure 4E** bottom) consistent with significant tumor progression (n= 10, *P* < 0.001, **Figure 4A** bottom).

Taken together, on Day2, pronounced increase of ADC and *K*^*trans*^ accompanying reduced tumor size revealed the pharmacodynamic effect on cancer cell and tumor microenvironment, providing biological insights at very early stage of KRASi therapy. mpMRI metrics exhibited consistent behavior on Day7, suggesting that they are robust. A responding tumor is likely to present with both elevated ADC and *K*^*trans*^ besides reduced tumor size.

When MRTX1133 treatment extended beyond Day7 in the survival study (**Figure 1B**), a median survival of 57 days in treated (n=20) vs. 22 days in CNTRL (n=20, *P* < 0.0001, Mantel-Cox test, **Supplementary Figure S5**) indicated a remarkable survival benefit. For treated mice, the baseline tumor size (or other MRI metrics) did not appear to predict survival since neither Pearson correlation nor regression analyses reached statistical significance.

### Responses of KRAS^G12D^ *vs* KRAS^G12C^ tumors to MRTX1133

Subcutaneous models grown from isogenic 4662-G12D and 4662-G12C cell line, respectively were enrolled for mpMRI study of therapeutic response on Day2 and euthanized after imaging (**Figure 1C**). The results are summarized in **Figure 5** and **Supplementary Figure S6**. Consistent with MRTX1133’s specificity, G12D subQ tumors exhibited a significant regression (**Figure 5A**) and an increase of K^trans^ (**Figure 5D**) while the on-target effect was confirmed by reduced pERK1/2 (n = 3, *P* < 0.05) and proliferation (n = 3, *P* < 0.001) compared to treated G12C tumors (**Figure 5E**). In contrast to increased ADC in response to KRASi therapy detected in KPC mice (Figure 2B), a majority of subQ G12D tumors exhibited a *reduction* in ADC on Day2 vs. baseline (P = NS, n=5, **Fig. 5B**, stared panel). Given ADC’s sensitivity to tumor cellularity, we suspect the unique feature of subQ G12D model might play a role: its fast growth was associated with the development of central necrosis in a fraction of tumors when tumor volume exceeded 50 mm^3^ whereas necrosis was usually absent in much slower growing G12C tumors. Although mice bearing visible necrotic tumors were not enrolled, the accumulation of inflammatory cells were observed in H&E sections of enrolled tumors (**Supplementary Figure S7**).

**Figure 5:**
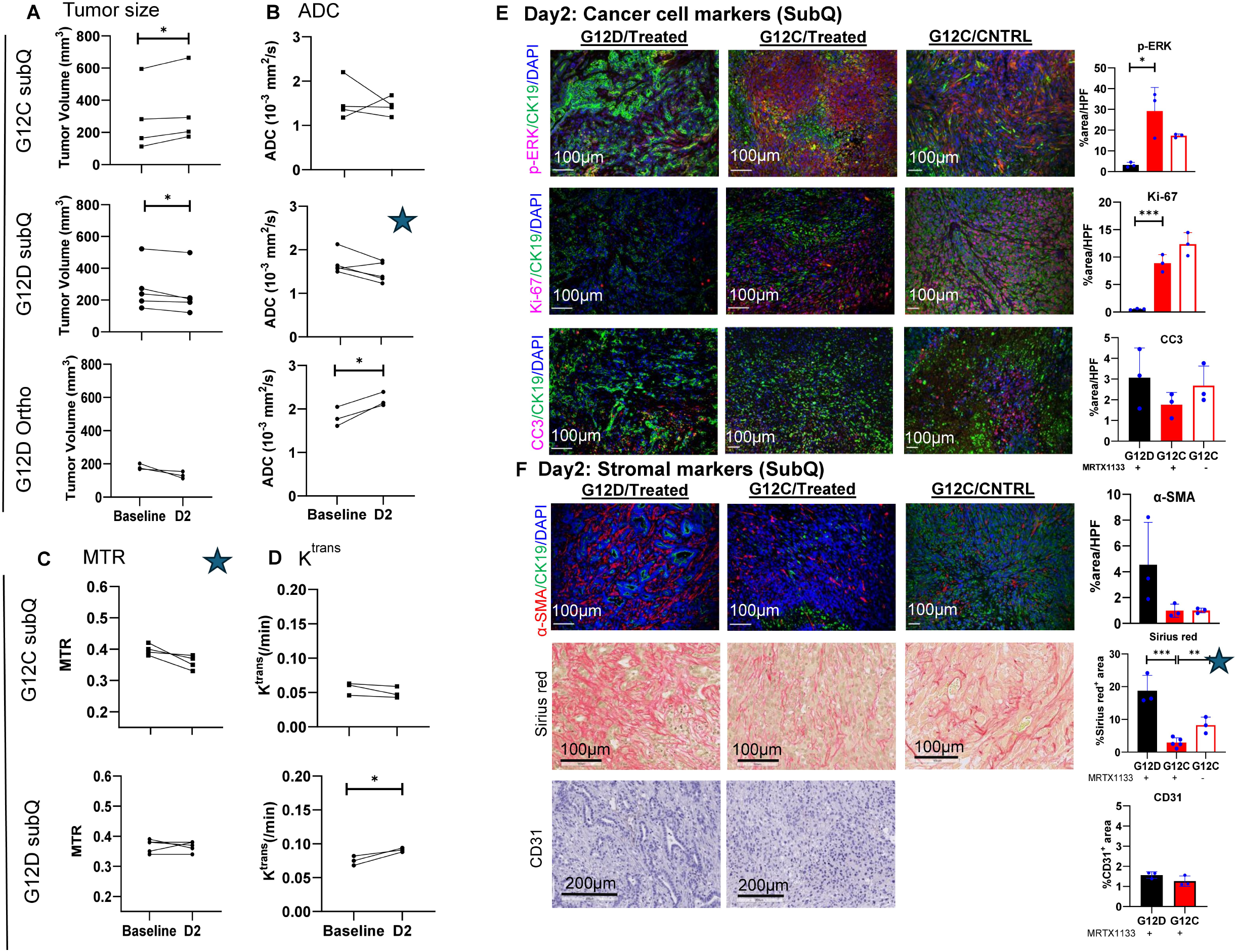
mpMRI and IHC of G12D vs. G12C mutated tumor responding to KRASi on Day2 (implant models). At baseline and Day2 of MRTX1133 treatment, tumor size (**A**), ADC (**B**), K^trans^ (**C**) and MTR (**D**) of subQ 4662-G12C and 4662-G12D tumors. Tumor size and ADC were also measured from orthotopic G12D tumors (A-B). (**E**) Cancer cell markers including p-ERK1/2, proliferation and death are assessed by double staining of p-ERK1/2 & CK19, Ki67 & CK19 and CC3 & CK19, respectively, in treated G12D vs. treated G12C vs. CNTRL G12C tumors. (**F**) Stromal markers of αSMA, Sirius red and CD31. αSMA-positive and CK19-negative cells were estimated in the plot. Sirius Red was stained in red and CD31-positive cells stained in brown color. Two-tail paired or unpaired t-test was applied when appropriate: \**P* <0.05, ** *P* <0.01, *** *P* <0.001. Starred panels represent unexpected results.

To test whether the unexpected ADC change is specific to subQ model, we inoculated 4662-G12D cells in the pancreas to produce orthotopic tumors and enrolled them when they grew to approximately 200 mm^3^. Increased ADC was observed in all treated mice (from 1.81±0.22 x10^−3^ mm^2^/s to 2.21±0.16 x10^−3^ mm^2^/s, n = 3, *P* < 0.05, **Figure 5B, ortho G12D panel**). Consistent to MRTX1133’s specificity to G12D mutation, the treated vs. CNTRL G12C tumors exhibited no difference in cancer cell markers (**Figure 5E**). However, a highly significant decrease in Sirius Red staining in treated G12C tumors (8.2±2.5 vs. 2.9±1.4 /CNTRL, n = 3-5, *P* < 0.01, **Figure 5F** stared panel) was detected. Interestingly, reduced Sirius Red was coincided with decreased tumor MTR values in G12C tumors after MRTX1133 treatment (n=4, *P* = 0.0534, **Figure 5C**, star-marked panel and **Supplementary Figure S6C**), suggesting that MTR could mark reduced collagen in G12C tumors to which MRTX1133 did not have on-target effects hence not introducing confounding MTR changes due to significant cancer cell death or cellularity changes.

In summary, results from isogenic tumors generated from 4662-G12D vs. 4662-G12C cell lines, respectively confirmed that the mpMRI can capture the drug’s specificity to G12D mutation while revealing a potential stromal effect on G12C tumors.

### mpMRI of tumors resistant to KRASi

Resistance to MRTX1133 treatment was confirmed in 17 out of 27 KPC mice enrolled (**Fig 1D**) while 10 died or were euthanized. Evolution of tumor size (**Figure 6A**) revealed that MRTX1133 treatment eradicated tumors in a subset of KPC mice temporarily, as their size fell beneath the radiographic threshold (dotted line) and became isointense with surrounding tissues on T2W MRI. However, tumor regrew shortly after treatment stoppage and eventually all surviving mice developed resistance defined as cancer outgrowth while on treatment (45). Tumor size, DWI and DCE metrics at the resistant stages were compared to the responding stage (**Figure 6B-D**) and to the baseline (**Supplementary Figure S8**). Tumor volume increased rapidly once resistance was established (R1/ 276±123 mm^3^/ n=17, R2 / 502±258 mm^3^/ n=14, R3 /688±481 mm^3^ / n=9, **Fig 6B**). ADC and KI values at R1/2/3 stages exhibited a significant reversal from those on Day2 and Day7 (*P* < 0.05 and *P* < 0.01, **Fig 6C**) while trending toward the baseline values (**Supplementary Figure S8B**). The reversal of mpMRI metrics was corroborated with the reversal of pERK1/2 (n=6, *P* < 0.01) and CC3 level in R3 tumors (n=6, *P* < 0.0001) compared to Day2 and/or Day7 (**Figure 6E**).

**Figure 6:**
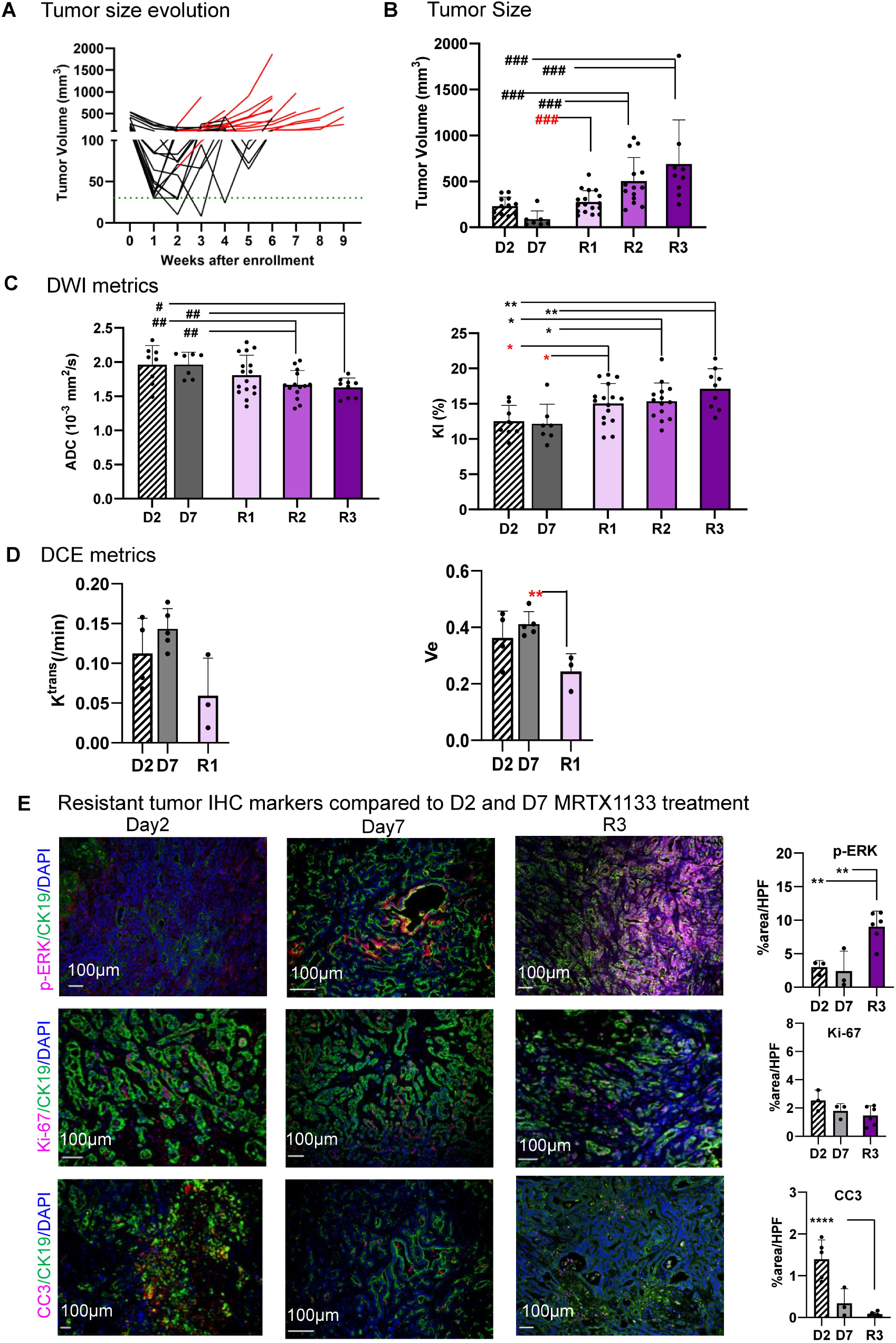
mpMRI and IHC markers of resistant tumors (KPC mice). (**A**) Tumor volume evolution during the development of resistance. Each line represents a mouse (red portion corresponds to resistant stage). (**B**) Tumor volume, (**C**) DWI metrics and (**D**) DCE metrics at resistant stages (R1, R2, R3) vs. responsive stages (Day2 and Day7). (**E**) Cancer cell markers of p-ERK1/2, Ki67 and CC3 of resistant tumor at R3 stage compared with those on Day2 and Day7. IHC micrographs of Day2 and Day7 are replicated from Figure 3 and 4, respectively. Mann-Whitney test was applied to data in (**B**,**C**) with ^#^ *P* < 0.05, ^##^ *P* < 0.01, ^###^ *P* < 0.001 while two-tail unpaired t-test applied to the rest with \**P* <0.05, ** *P* <0.01, *** *P* <0.001, **** *P* <0.0001. Red * or # marks the mpMRI metrics at R1 stage whose values differed significantly from responding stages (Day2 or Day7).

Assessing whether Day7 mpMRI can predict resistance development, we found that tumor size did not correlate with the time to R1 occurrence (Pearson correlation coefficient, r = 0.011, P = 0.97, n=17). However, a trend of negative correlation was observed between ADC and time to R1 (r = −0.83, P = 0.08, n= 5). Despite relatively small sample size, the analysis of ADC’s predictive power could help generate hypothesis for future studies.

In summary, upon development of resistance to KRASi, mpMRI metrics exhibited a reversal of values from responsive stages, revealing increased heterogeneity of diffusion (KI ++), reduced extracellular & extravascular volume fraction (V_e_ --) consistent with increased cellularity (ADC --) and disease progression (tumor volume +++).

## Discussion and Conclusion

We present the first multi-parameter MRI study to evaluate pancreatic cancer responses to KRAS inhibitor therapy. In this study, we examined clinically applied MRI metrics including tumor volume, DWI (ADC, KI), DCE (K^trans^, V_e_), MTR to evaluate responses of PDAC tumors to KRASi (MRTX1133) treatment on Day2, Day7 and after prolonged treatment that leads to acquired resistance. While tumor size provides an objective measure of KRASi effect and a reference for the development of drug resistance, ADC and K^trans^ offer insights of cell death, reduced cellularity and significant stromal changes underlying tumor size reduction. On Day2 post treatment, significant increases in ADC are indicative of cell death that is confirmed by CC3 staining and pronounced reduction of pErk1/2 level in cancer cells, a marker of MRTX1133 engagement to its target. Maps of mpMRI metrics on Day2 revealed remarkable intratumor heterogeneity compared to baseline induced by KRASi treatment (**Figure 2**) corroborated by histopathology (**Supplementary Figure S3**). On Day7, tumor ADC values remain elevated, consistent with sustained pERK1/2 suppression. KI and V_e_ (EES) become significant on Day7 primarily due to deepened therapeutic effects. Our data demonstrate a strong positive correlation between ADC vs. CC3 and an inverse correlation between ADC vs. cellularity (**Figure 3C,D**), supporting the hypothesis that a sharp increase of ADC early after KRASi treatment marks cell death and/or reduced cellularity in responding PDAC tumors. In contrast to autochthonous or orthotopic model, influx of inflammatory cells occurred in subQ G12D tumors masked the ADC increase induced by KRASi (**Figure 5B star panel**).

Besides the effects on cancer cells, KRASi induces rapid and pronounced changes in stroma markers. Significant and consistent increase of K^trans^ was detected on Day2 and Day7 and accompanied by reduction of microvascular density (**Figure 3E**). Hence KRASi treatment may have vascular “pruning” effect that reduces angiogenesis, leading to vascular “normalization” and increased blood perfusion. Depletion of matrix hyaluronan by hyaluronidase (PEGPH20) also led to increased K^trans^ in KPC tumors without changes of microvascular density (32) via a different mechanism, where capillaries recover from collapsed state and resume perfusion when tumor interstitial fluid pressure is reduced after PEGPH20 treatment (46). While K^trans^ metric obtained from current kinetic modeling is robust, perfusion and permeability contribution to the K^trans^ metric is unseparated. We strive to resolve this issue by more sophisticated kinetic model (47) in future studies.

Our observation that MTR does not respond to increased collagen content in KPC or 4662-KRAS^G12D^ tumors after MRTX1133 treatment suggest that the MTR value is determined by two opposing effects: an increase in bound water pool due to increased collagen deposition *versus* reduced bound water upon cell death, both mediated by KRASi. Hence, in treatments that induce substantial cell death, MTR would *not* be a suitable marker for collagen level. However, in 4662-KRAS^G12C^ tumors where the treatment did not induce significant cell death or cellularity change, MTR seemed able to mark the change of extracellular matrix collagen content.

Compared to MRTX1133, which binds to KRAS protein in GDP-bound (OFF) state, RASi specific to KRAS in GTP-bound (ON) state, to HRAS, NRAS or wild type RAS have been developed (7–10) as well as other modalities of direct RAS inhibition via binder protein, siRNA and CAR-T cells have been developed (11,12). Regardless of modalities that are used to inhibit RAS, the downstream effects of cancer cell death and stromal reprogramming upon RAS pathway blockade are likely to be similar therefore the mpMRI is expected to have general applicability for all forms of RAS inhibition.

Along with technical advancements that have greatly improved the DWI and DCE performance of clinical MRI system, international consensus is being developed for standardization of acquisition and analysis protocols for DWI and DCE across clinical MRI platforms (48–50). The coordinated efforts from academia, professional societies and industry aiming to enhance robustness and repeatability of MRI metrics in multi-center clinical studies are likely to facilitate the clinical translation of these imaging tools to evaluate patient responses to KRASi therapy.

## Supporting information

list of abbreviations

## Acknowledgement

This study was supported by U24-CA231858 (Penn Pancreatic Cancer Imaging Resource) and R01CA266285. We want to thank Miguel Romanello Joaquim for technical assistance and are grateful for the support of institutional resources, especially the Mouse Hospital of Penn Pancreatic Cancer Research Center and the Small Animal Imaging Facility of Radiology Department, University of Pennsylvania.

