## Supplementary figures for "Multimetric MRI Captures Early Response and Acquired Resistance of Pancreatic Cancer to KRAS Inhibitor Therapy"

**Supplementary Figure S1**: **mpMRI metrics and corresponding parameter maps of the tumor. (A)** Tumor size is obtained from T2-weighted (T2W) MR images. Region Of Interest defined for tumor, spinal muscle, phantom-1 and phantom-2 (10% and 40% Polyvinylpyrrolidone, respectively) are marked. The phantoms serve as references for ADC as their ADC values were quantified by independent method and bookend the ADC range of KPC tumors (38). **(B)** ADC and KI metric are calculated from diffusion weighted images of five b values (detailed in Methods). **(C)** DCE metrics ($K^{trans}$, $V_e$ and $\tau_i$) are derived from kinetic modeling of DCE series along with T10 map and group arterial input function (41) with B1 correction as we described previously (40). **(D)** MTR metric and $MTR_{bin}$ histogram are also detailed in Methods.

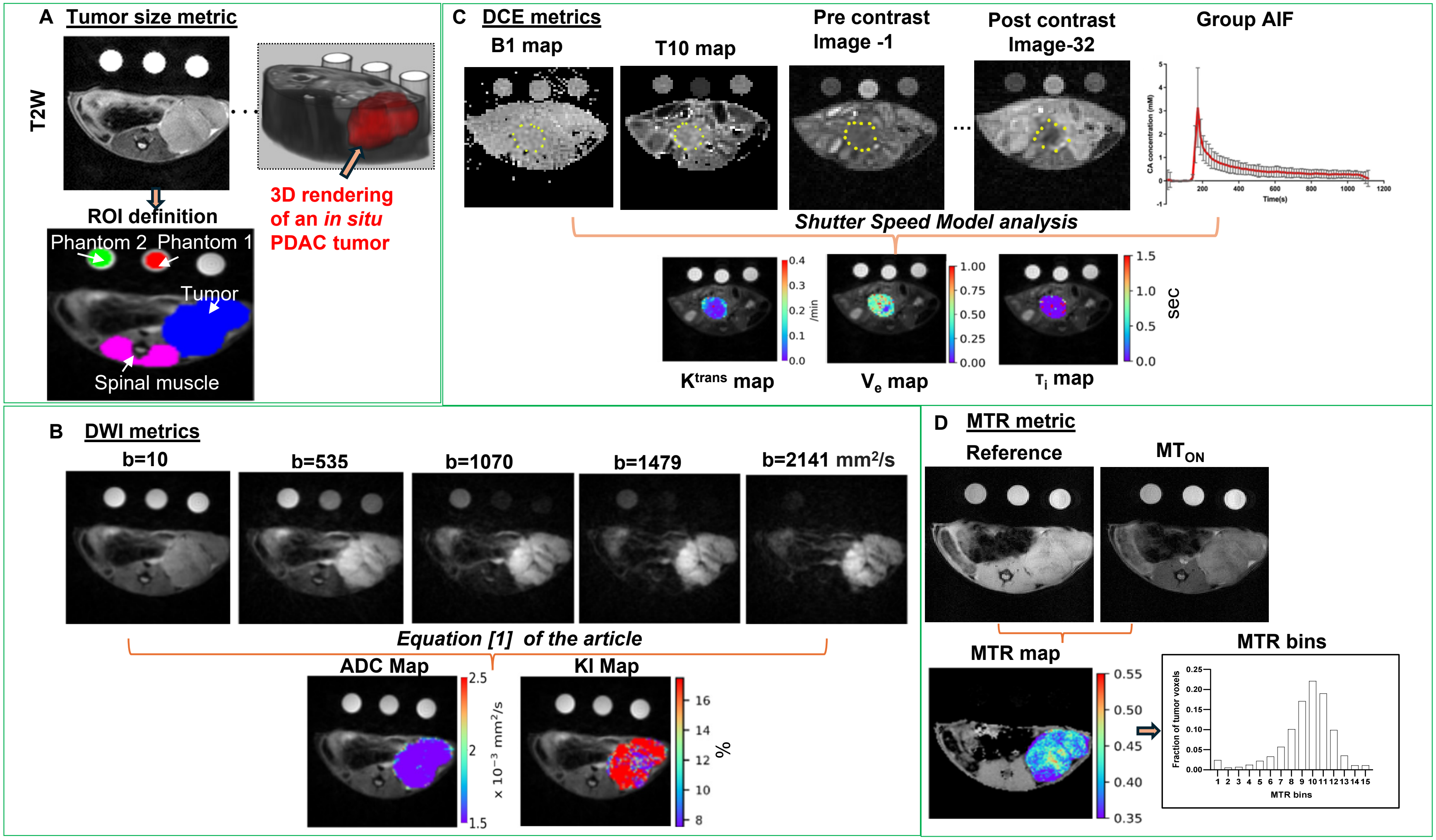

**Supplementary Figure S2: Tumor collagen content from CNTRL and MRTX1133 treated KPC tumor** Tumors were collected from CNTRL (n=5) and treated mice on Day2 (n=8) and Day7 (n=4). Tumor collagen content was estimated by biochemical assay (see Methods). Two-tail, unpaired t-test was used to compared treated vs. CNTRL. ** *P* <0.01, *** *P* <0.001

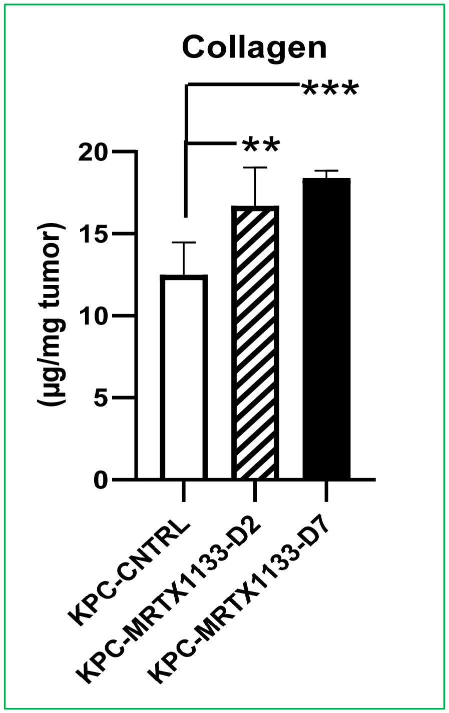

**Supplementary Figure S3: Day2 IHC revealed intratumor heterogeneities after KRASi treatment.** H&E **(A)** and Sirius Red **(B)** micrographs of whole tumor section with selected regions labeled as box1-4. Both whole section and regional analysis of cellularity and collagen content was conducted by a GI pathologist (EEF) as detailed in Methods. Representative T2W images, ADC, KI and MTR maps of this tumor (from KPC mouse #30292) on Day2 and baseline are shown in **Figure 2A-D**.

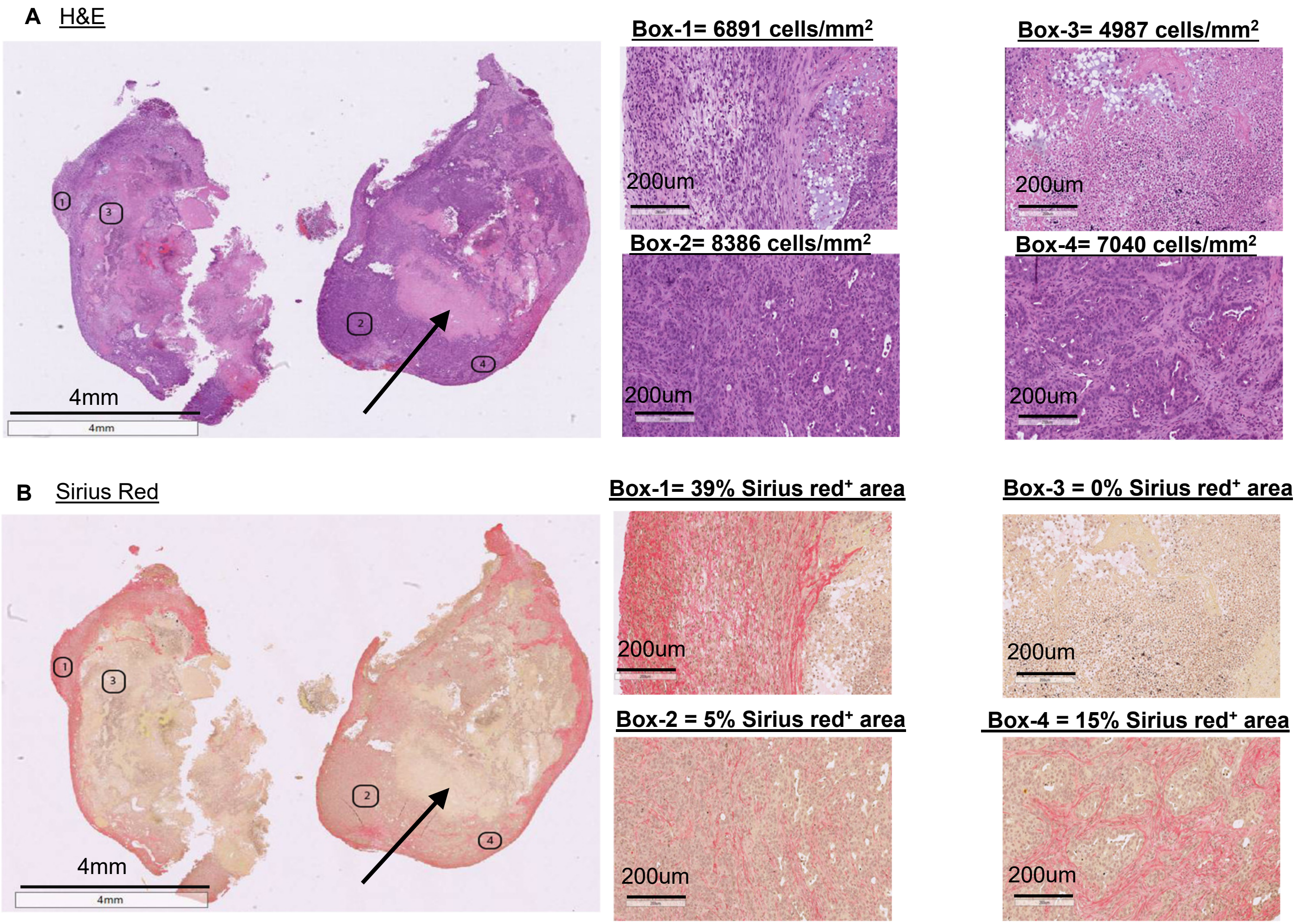

**Supplementary Figure S4: Day7 mpMRI of treated vs. CNTRL mice.** Three slices from the same tumor are shown for each metric acquired on Day7 and baseline. Tumor size and DW metrics (**A, B, C**) were from the same mouse whereas DCE metrics **(D** and **E)** from a different mouse in the treated group while all metrics were from the same mouse in CNTRL group.

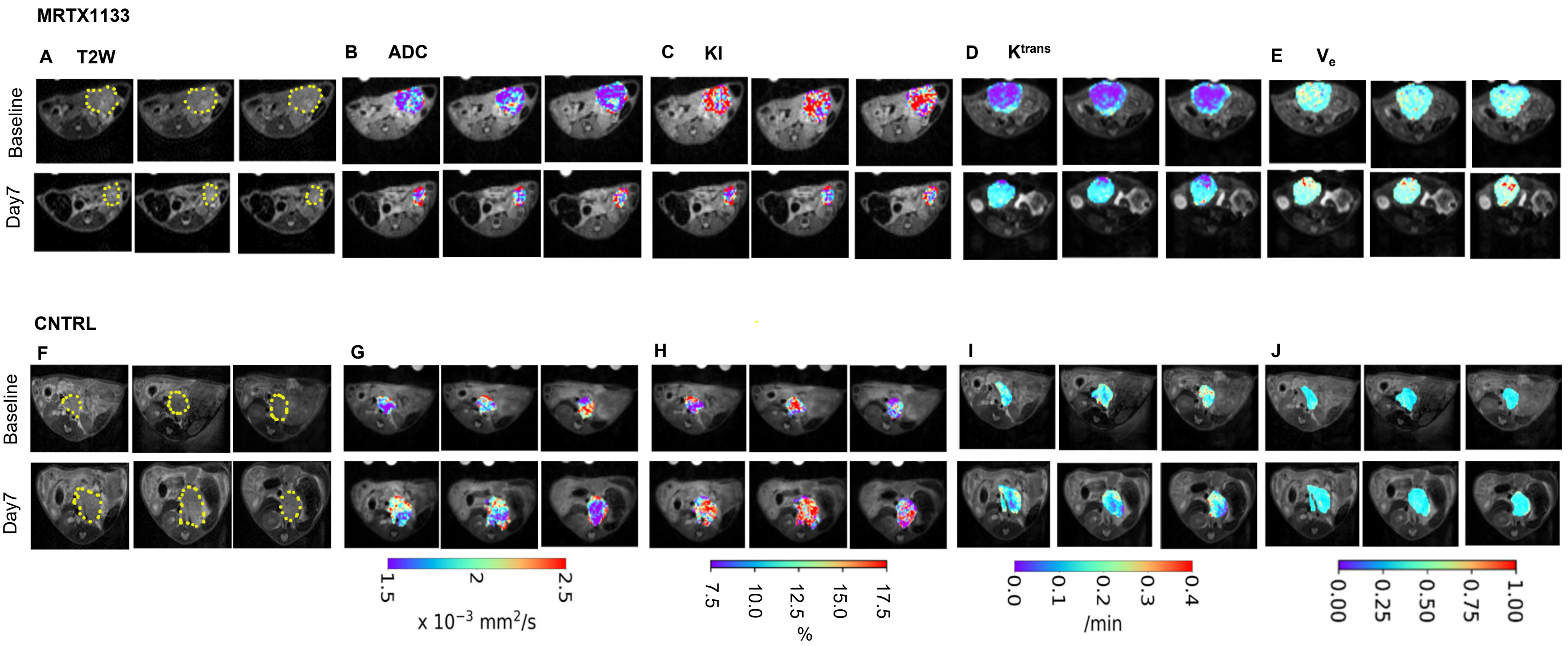

**Supplementary Figure S5: Kaplan-Meier curve of treated and CNTRL KPC mice.** Median survival is 57 days for MRTX1133 treated group (n=20) and 22 days for CNTRL group (n=20). ****$P$ < 0.0001, Mantel-Cox nonparametric test.

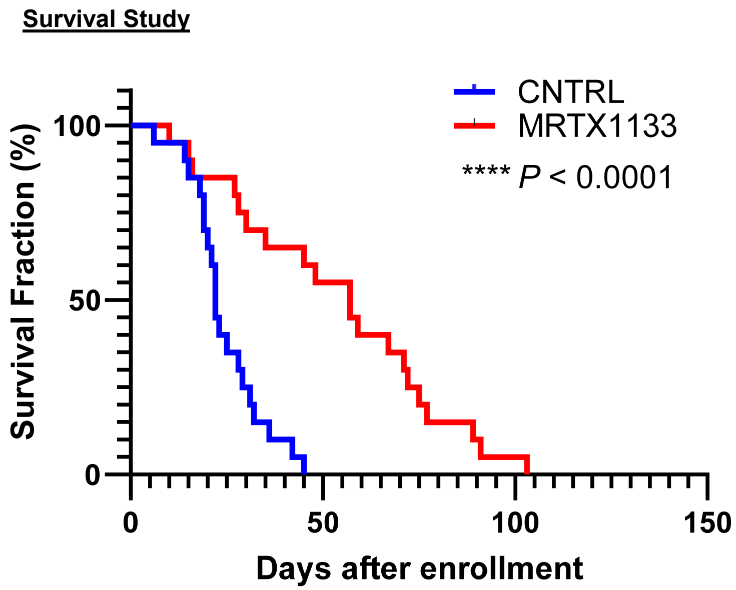

**Supplementary Figure S6: Day2 mpMRI of 4662-G12C and 4662-G12D subQ tumors and 4662-G12D orthotopic tumors.** Tumor Region Of Interest are marked on T2W images with dotted lines (**A**), maps of ADC (**B**), MTR (**C**) and $K^{trans}$ (**D**) at baseline and on Day2 are overlaid on T2W MRI. Four image slices are shown for the same tumor at each time point.

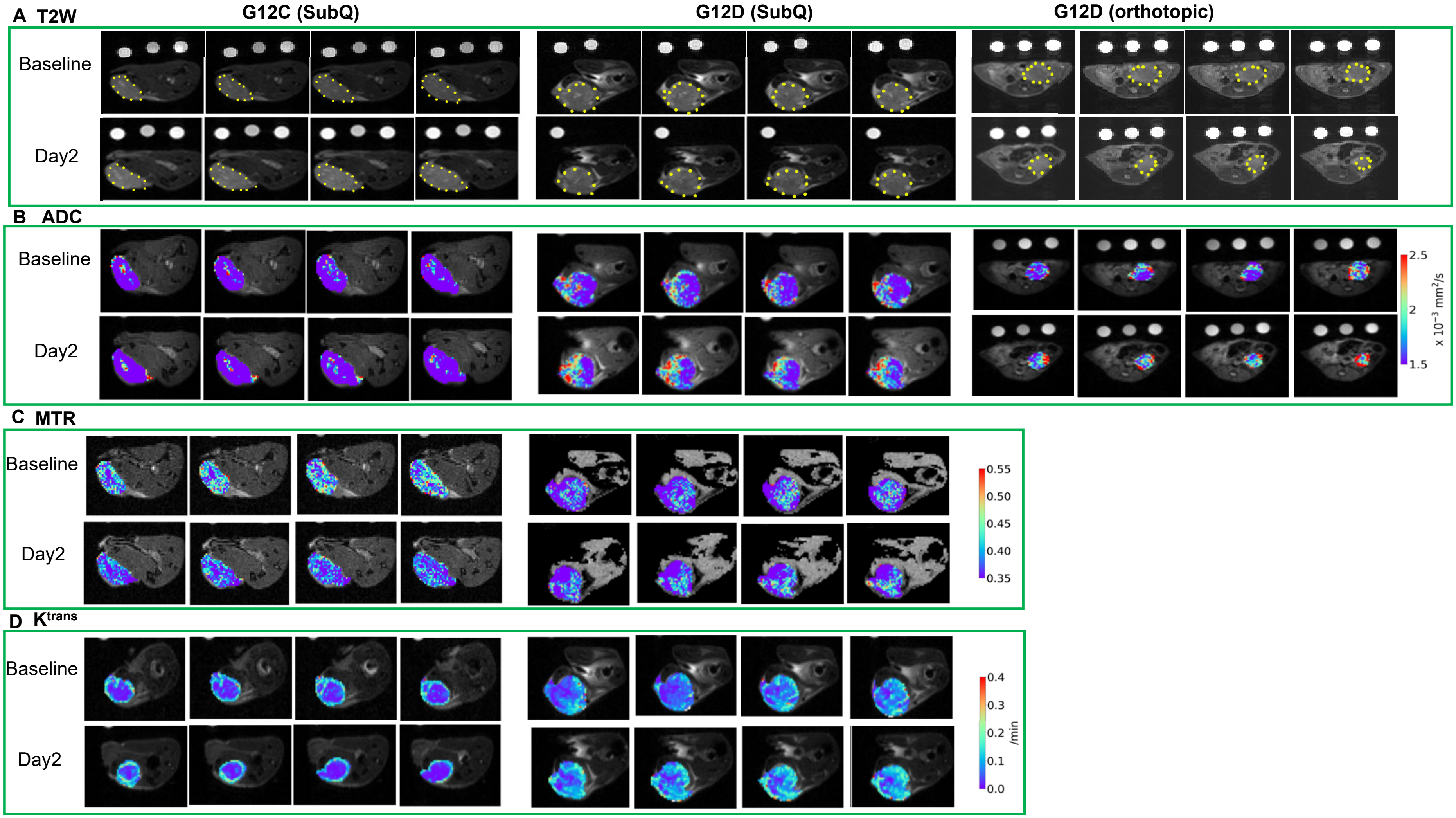

**Supplementary Figure S7: H&E section of a 4662-G12D subcutaneous tumor. A magnified view from a viable tumor region (black box) is shown where many mononuclear cells and occasional neutrophils are observed among the adenocarcinoma cells. For clarity, selected inflammatory cells are marked by white arrows.**

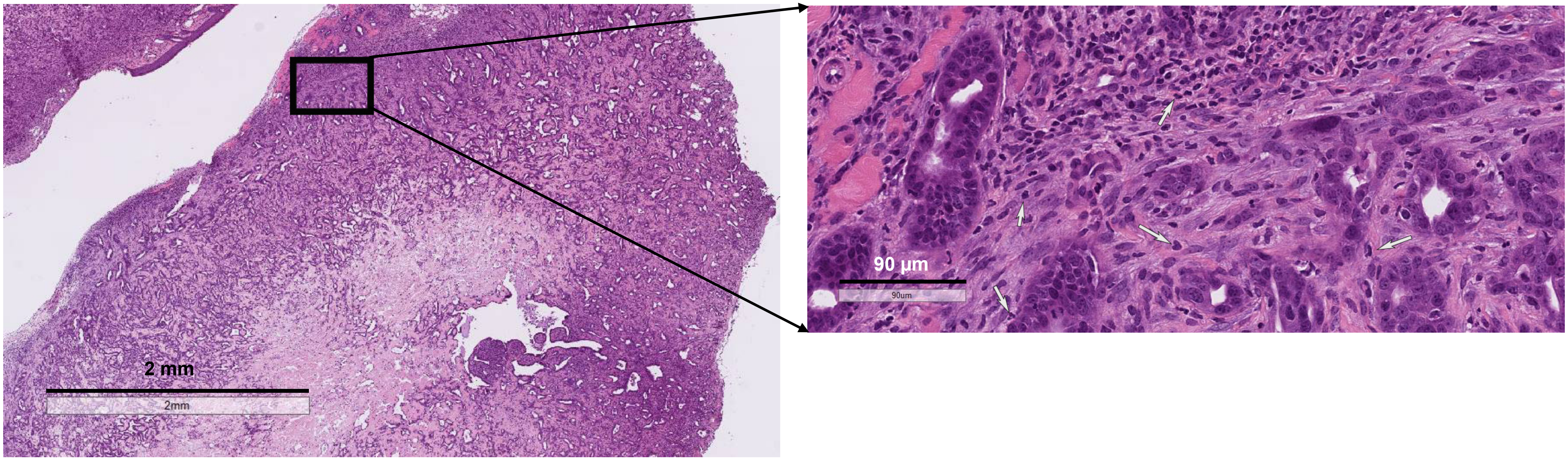

**Supplementary Figure S8: mpMRI markers at resistant stages vs. baseline in KPC mice**. Tumor size (**A**), DWI metrics (**B**), $MTR_{bins}$ (**C**) at R1, R2, R3 stage respectively are compared with baseline while DCE metrics of resistant tumor was only obtained at R1 to compare with baseline (**D**). Two-tail unpaired t-test was conducted. **P* <0.05, ** *P* <0.01. Red star (*) marks the mpMRI metric whose values differed significantly at early resistant stage (R1) vs. baseline.

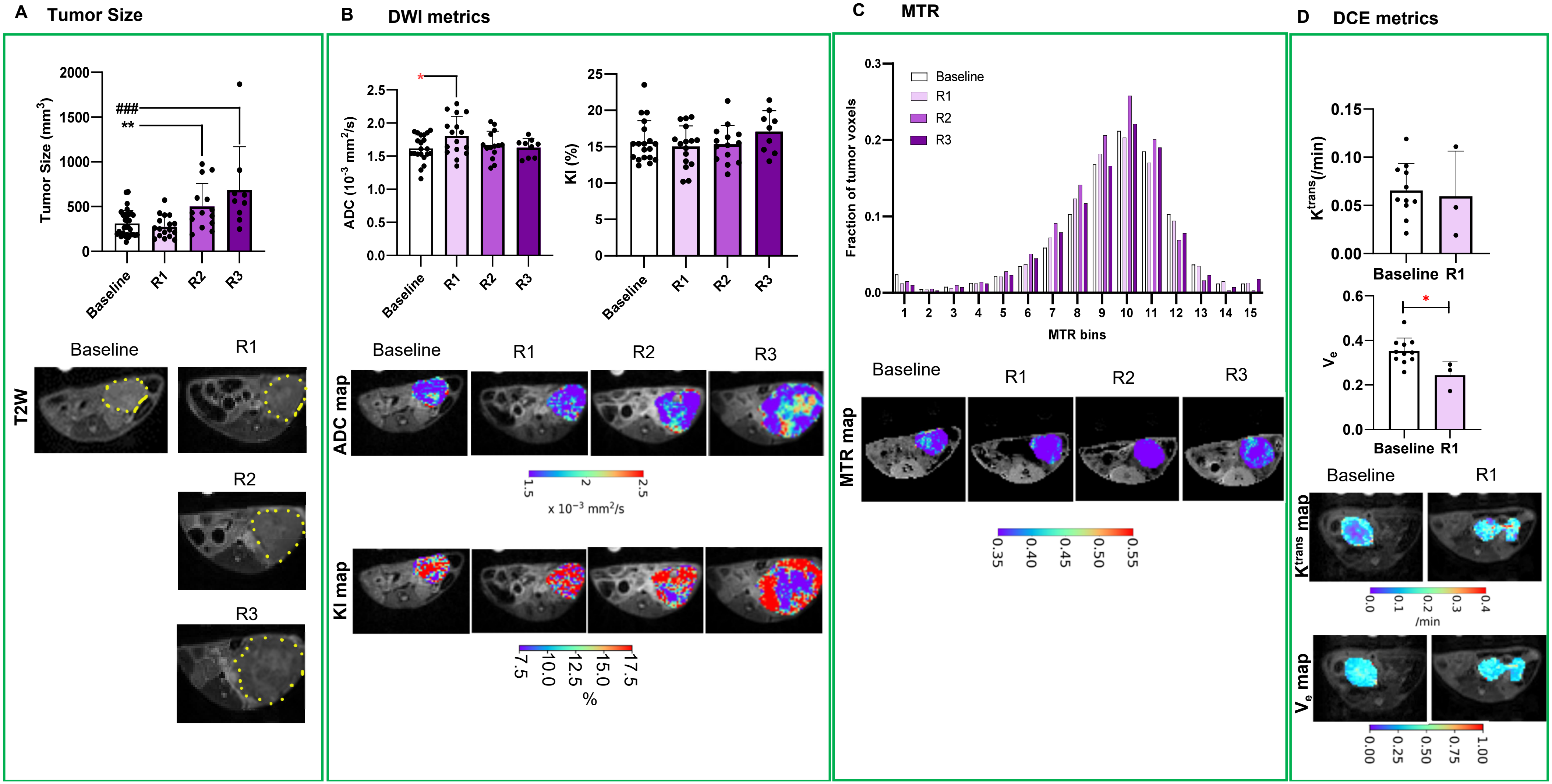
