## Supplementary material for "Multimetric MRI Captures Early Response and Acquired Resistance of Pancreatic Cancer to KRAS Inhibitor Therapy": list of abbreviations

**Supplementary List of Abbreviations**

| | |
|---|---|
| **ADC** | Apparent Diffusion Coefficient |
| **CNTRL** | Control (Group) |
| **CC3** | Cleaved Caspase-3 |
| **DCE** | Dynamic Contrast-Enhanced MRI |
| **DWI** | Diffusion weighted MRI |
| **IHC** | Immunohistochemistry |
| **KRASi** | KRAS Inhibitor |
| **KPC** | *LSL-Kras*$^{G12D/+}$; *LSL-Trp53*$^{R172H/+}$; *Pdx-1-Cre* mice |
| **KI** | Kurtosis index as defined in equation [1] of Methods |
| **$K^{trans}$** | Rate constant of transferring unit volume of contrast agent from capillaries to interstitial space per minute |
| **mpMRI** | Multiparametric MRI |
| **MTR** | Magnetization Transfer Ratio |
| **PDAC** | Pancreatic Ductal Adenocarcinoma |
| **TME** | Tumor Microenvironment |
| **$V_e$** | Extracellular and extravascular volume fraction |
