## Supplementary Table S1 for "Multimetric MRI Captures Early Response and Acquired Resistance of Pancreatic Cancer to KRAS Inhibitor Therapy"

**Supplementary Table S1: Catalog numbers for monoclonal antibodies and dilution schemes used in immunofluorescence.**

| Antibody | Supplier | Catalog Number | IF dilution |
|---|---|---|---|
| p-ERK1/2 | Cell Signaling | 4370 | 1:100 |
| Alpha-smooth muscle actin (aSMA) | Sigma-Aldrich | A2547 | 1:1000 |
| Cleaved caspase 3 (CC3) | Cell Signaling | 9661 | 1:100 |
| Ki-67 | eBioscience | 14-5698-82 | 1:100 |
| CK19 (TROMA-III) | Iowa Developmental Hybridoma Bank | AB-2133570 | 1:200 |
| Donkey anti-Rat IgG (H+L) Highly Cross-Adsorbed Secondary Antibody, Alexa Fluor™ 488 | Invitrogen | A-21208 | 1:250 |
| Donkey anti-Mouse IgG (H+L) Highly Cross-Adsorbed Secondary Antibody, Alexa Fluor™ 647 | Invitrogen | A-31571 | 1:250 |
| Donkey anti-Rabbit IgG (H+L) Highly Cross-Adsorbed Secondary Antibody, Alexa Fluor™ 594 | Invitrogen | A-21207 | 1:250 |
| Donkey anti-Rabbit IgG (H+L) Highly Cross-Adsorbed Secondary Antibody, Alexa Fluor™ 488 | Invitrogen | A-21206 | 1:250 |
| Donkey anti-Rat IgG (H+L) Highly Cross-Adsorbed Secondary Antibody, Alexa Fluor™ 594 | Invitrogen | A-21209 | 1:250 |
